# Variability of Head Motion Reveals Development of Spatial Attention

**DOI:** 10.64898/2026.09.23.753739

**Authors:** Malte Wöstmann, Dunja Kunke, Nicole Wetzel

**Author notes:** Corresponding author, Malte Wöstmann, Department of Psychology, University of Lübeck, Lübeck, Germany Maria-Goeppert Strasse 9a, 23562 Lübeck.

## Abstract

Selective attention, the ability to focus on relevant information while ignoring task-irrelevant information, is crucial for learning. Selective attention has largely been studied in motionless individuals. However, body movements are intrinsic to cognition and particularly frequent in children. To test whether movements explain successful attention control throughout childhood, we tracked head rotations while younger children (6–8 years, n = 25), older children (9–10 years, n = 20) and adults (18–30 years; n = 21) identified a spatial target sound under distraction. Distractor reports decreased with age, confirming a developmental increase in attention control. Importantly, a dissociation of two distinct types of movement emerged: Task-dependent head rotation to lateral targets was similar across age groups. Contrary, task-independent movements before and early during a trial decreased with age and predicted attention accuracy. Thus, movements unrelated to the attention goal do not reflect noise but signify attentional control across development.

## Introduction

Humans perform a rich repertoire of movements at any time, ranging from automatic physiological movements (e.g., related to respiration or heartbeat), over postural sway to voluntary movements of individual body parts. Traditionally, experimental psychology has tried to study cognitive functions independent of body movement, by fixating study participants (e.g. using a chin rest) or by rejecting movement-contaminated data (for reviews, see Ladouce et al., 2017; Stangl et al., 2023). However, embodied cognition presumes that cognitive processes are shaped by the body’s morphology, sensorimotor capacities and interactions with the environment (Clark, 1999; Wilson, 2002). The concept of active sensing implies that the human mind is no passive observer but employs active motor routines to control sensory inflow (Morillon et al., 2015; Schroeder et al., 2010). More recently, propelled by evidence from technologies that precisely measure bodily states, the view that cognition originates from interaction of brain and body gains renewed support (Criscuolo et al., 2022; Mathis et al., 2018; Pereira et al., 2020; Stringer et al., 2019).

A core cognitive function is attention, which allows to focus on relevant information and to prevent distraction (Desimone & Duncan, 1995; Noonan et al., 2018; Wöstmann et al., 2022). Influential theories posit a strong involvement of motor activity in attention, assuming that shifts of attention base on the preparation of orienting movements (Rizzolatti et al., 1987) and that attention emerges from the selection of competing sensorimotor representations (Cisek, 2007). As humans cannot move or close their ears, auditory attention involves head movements to modulate the sensory sampling of competing sounds (Grange & Culling, 2016; Kondo et al., 2012). Recently, we found that adult listeners perform prominent head rotations in the direction of task-relevant sounds when movement is permitted, as well as miniature head rotations when they are asked not to move (Wöstmann & Obleser, 2026). This suggests that auditory spatial attention is tightly coupled to the preparation and execution of motor actions. However, we currently lack an understanding of whether and how variability of motor activity shapes successful attention to relevant sound.

Childhood development is an important model for investigating the relation of movement and attention. The number of limb movements per hour varies across development, showing an inverted U-shaped pattern with a peak in middle childhood at the age of ∼7–9 years (Eaton et al., 2001). At the same time, distractibility by irrelevant sounds decreases (Hoyer et al., 2021; Wetzel et al., 2019), suggesting increasing attentional control, especially from ∼6–10 years of age. Motor activity in children has been linked to different aspects of cognition, such as improved learning (Agostinho et al., 2015), saving cognitive resources (Ping & Goldin-Meadow, 2010), or higher working memory demands (Rapport et al., 2009). However, despite research showing that physical exercise or general fitness relate to children’s attention performance (Janssen et al., 2014; Reigal et al., 2019), it is to our knowledge unknown whether movement patterns in environments with competing sources of relevant and irrelevant information relate to the development of attention. Understanding this relationship carries substantial societal relevance, as orienting movements toward target sounds or away from distractors may help counteract the negative effects of acoustic noise (e.g., in classrooms), which is known to impair children’s academic attainments (Caviola et al., 2021; Dockrell & Shield, 2006).

Research on movement during cognitive tasks has traditionally focused on actions instructed by the task itself, such as button presses or eye movements toward a target. However, a growing body of work shows that task-independent movements (e.g., fidgeting, grooming, postural shifts, and other unprompted actions) are pervasive during task performance and dominate trial-by-trial neural variability in mice (Musall et al., 2019). Critically, these task-independent movements are not simply noise to be filtered out. It has been shown that although the overall frequency of movements does not change as mice shift from engaged to disengaged states, movements become more erratically timed and idiosyncratic during disengagement, and this shift in movement patterning predicts task performance (Yin et al., 2025). This suggests that task-independent movement is not a peripheral nuisance variable but might constitute a behavioral signature of the underlying cognitive state in humans as well.

Here, we tested two closely related hypotheses regarding the contribution of body movements to auditory spatial attention from a developmental perspective. First, we predicted that elementary-school children (6–10 years) would increase systematic motor action (rotation of the head) in service of auditory selective attention, demonstrating the development of embodied cognition (Loeffler et al., 2016). Second, we hypothesized that isolating task-independent from task-dependent movements would uncover relationships between head movements and selective attention and how they change during childhood and adolescence. We used a gamified version of an established auditory spatial attention paradigm wherein a lateral and a frontal sound competed for attention (Wöstmann et al., 2019, 2025; Wöstmann & Obleser, 2026). A head-mounted gyroscope continuously tracked patterns of head rotation. Our results demonstrate that task-dependent head rotation is largely stable across childhood development whereas task-independent head rotation decreases during development and predicts the success of attentional selection.

## Method

### Participants

Adults, n = 21 (*M*_age_ = 24;6 (years; months), range 18;4-30;3, 10 females, 1 left-handed, all native German), n = 25 native German younger children (*M*_age_ = 7;8, range 6;0-8;11, 14 females, 4 left-handed), and n = 20 older native German children (*M*_age_ = 9;8, range 9;0-10;11, 13 females, none left-handed) were included in the final sample. Handedness was measured with an abbreviated German version of the Oldfield Handedness Inventory (Oldfield, 1971). Data of five additional children were recorded but excluded from the final data analysis due to technical issues resulting in incomplete datasets: 3 younger children; task performance < 20% correct: 1 older child; no convergence of staircase procedure to titrate SNR: 1 younger child. The local ethics committee of the Otto-von-Guericke University Magdeburg approved all experimental procedures (No.164/24). The rationale for the sample size was to use a factorial design (with three age groups) to obtain effects with a partial eta-squared (*η^2^_p_*) ≥ .15 with 80% power, which required at least 19 participants per age group.

The participating families were recruited from local public and private schools. Parents of the participating children and adults gave written informed consent and the children agreed verbally. Adult participants and parents confirmed that none of the exclusion criteria applied to them or to their children (disabilities in hearing or eyesight that could not be corrected by glasses, neurological disorders or medication that influence the central nervous system). Children received a voucher from a local toy store after the experiment. Adult participants received €10/hour.

### Stimulus material

We used pictures of animals (penguin & frog) as spatial attention cues. All visual stimuli were presented on a grey background (RGB: 128, 128, 128). The screen resolution was 1366 x 768 px, with spatial attention cues covering approximately 372 x 372 px (Fig. 1). The position of the animal’s pupil (left, right, front) indicated the to-be-attended loudspeaker location. Visual stimuli were presented on the screen of a laptop (Dell, running Windows 10 Pro) with a screen size of 15.6".

**Figure 1.**
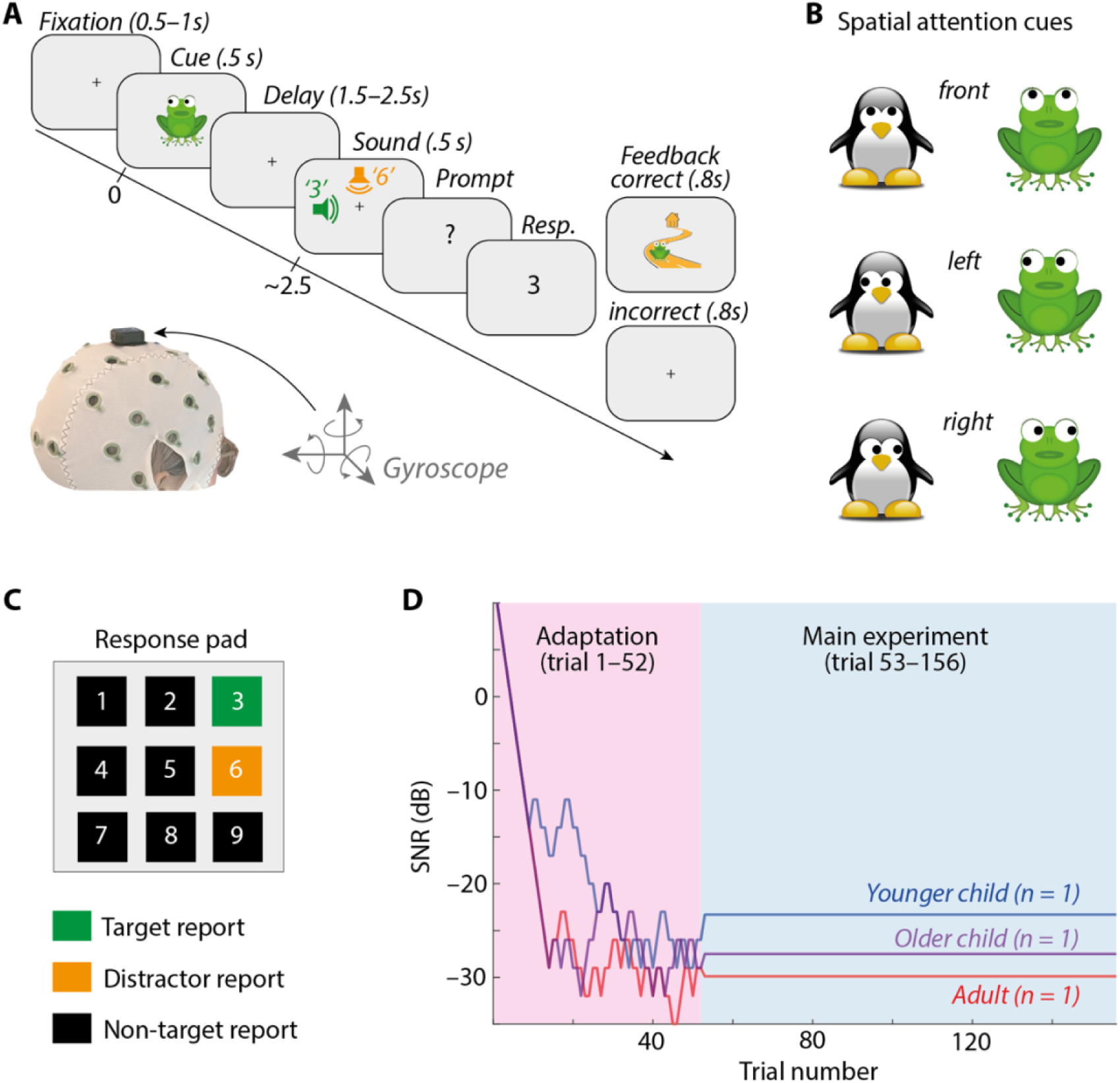
(**A**) Top: design of auditory spatial attention task. Bottom: placement of the gyroscope on a participant’s head. (**B**) Pictures of animals were used as spatial attention cues (penguin, frog), directing their gaze to the to-be-attended sound location (front, left, right). (**C**) Response pad. Colors of buttons indicate different response options for a trial with the target number “3” and the distractor “6”. Buttons were not colored in the actual experiment. (**D**) Individual adaptation of the SNR of acoustic stimuli across time (adaptation blocks 1&2, main experiment blocks 3–6) for three representative participants.

Auditory target and distractor stimuli were spoken numbers (1–9, same female voice), each adjusted to a duration of 0.5s (Wöstmann et al., 2025). The cover story of the experimental task instructed participants to help the animal find the way back home. Feedback was provided in correct trials only, showing an image of the animal on its way home.

### Procedure

We used an adapted version of a previously established auditory spatial attention paradigm (Wöstmann et al., 2019, 2025; Wöstmann & Obleser, 2026). One loudspeaker was always placed in front of participants (0° azimuth; 0° elevation; distance: 63 cm), the other loudspeaker was placed on the left or right (–90° or +90° azimuth; 0° elevation; distance: 63 cm) alternating in a block-wise fashion. Given these two setups and two possible spatial cueing conditions (cue pointing to frontal versus lateral loudspeaker), four experimental conditions were implemented: select-left (speakers: front and left; spatial cue: left), select-right (speakers: front and right; spatial cue: right), suppress-left (speakers: front and left; spatial cue: front), suppress-right (speakers: front and right; spatial cue: front).

The experimenter read the instructions to the children while the adult participants read the instructions themselves. Participants were instructed that the experimental task was to help the animals find their way back home. Accordingly, the cued target number was told to be uttered by the respective animal’s mother, who tried to tell the animal how many steps to proceed in a certain direction. The participant’s task was to help the animal understand the target number by selecting it on the response pad at the end of a trial. Participants were additionally instructed that they were permitted to move and turn their head to facilitate identification of the target number.

Before the beginning of the actual experiment, participants performed five instructed training trials together with the experimenter. The training served the purpose to make sure participants understood the task. For the first two trials, spatial cues were shown on the screen and target numbers were presented without distractors. The last three training trials contained targets and distractors, as in the actual experiment. After successful completion of training trials, the actual experimental procedure was started.

Each trial of the experiment started with the presentation of the spatial cue (animal looking to left, right, or front) for 0.5s (Fig. 1). Next, a fixation cross was shown in the center of the screen for a variable delay period (randomly jittered from 1.5–2.5s in steps of 0.01s). After the delay, two different spoken numbers were presented simultaneously from two positions for 0.5s (front-left or front-right). The number presented at the cued location was the target, the other number was the distractor. Next, a question mark was shown, and participants used an external number pad to report the target number. The question mark was replaced by the participant’s response as soon as a button on the number pad was pressed. Feedback was then presented for 0.8s (correct: positive feedback screen, incorrect: fixation cross).

Each participant performed a total of 156 trials, divided in 6 blocks à 26 trials. The position of the lateral loudspeaker (left vs. right) changed before the beginning of every new block. The first two blocks implemented a staircase procedure to adjust overall performance to ∼71% task accuracy (Levitt, 1971). Of the first two blocks, one used the penguin and the other the frog as the spatial cue, balanced across participants. The initial signal-to-noise ratio (SNR) of auditory targets versus distractors was set to +10dB. An adaptive 1-up, 2-down procedure enhanced the SNR after one incorrect trial and lowered it after two consecutive correct trials in steps of 3dB (by varying the intensity of the target number while keeping the distractor’s intensity constant). In the end of the second block, the individually titrated SNR was calculated as the average SNR on the last 5 trials of the two blocks. The SNR in the four remaining blocks was set to the SNR titrated in the first two blocks. Note that both behavioral responses and head rotations recorded during the first two blocks were not considered in the data analysis. The experiment was implemented using the Psychophysics Toolbox (Brainard, 1997) and the Palamedes Toolbox (Prins & Kingdom, 2018) for Matlab (R 2020a).

After titration of the SNR, participants performed the remaining 4 blocks (104 trials), resulting in 26 trials per condition in the 2 (attention: selection vs. suppression) x side (left vs. right) design, with *attentional selection* referring to trials with a lateral target and frontal distractor and *attentional suppression* referring to trials with a lateral distractor and frontal target. One cue animal (penguin, frog) was used in blocks 3&4 and the other in blocks 5&6, with the order of cue animals balanced between participants. In-between every two blocks, participants could take a short break of 2 to 3 minutes.

### Gyroscopic tracking of head rotations

Head movement was recorded using a small wireless (Bluetooth) inertial sensor (Witmotion, ShenZhen, China). The effective sampling rate of the sensor, which varied slightly around 20Hz, was linearly resampled to 20Hz. To standardize the placement of the sensor, an EEG cap (Brainvision actiCAP with SNAP holders) was placed on the participant’s head and the sensor was placed at the vertex position (electrode position Cz). Head rotation was measured in angular velocity (°/s) around three orthogonal rotational axes (yaw, pitch, roll). Additionally, acceleration along three axes was assessed, but not analyzed for the purpose of the present study. Note that although head rotation was also recorded during the SNR adaptation (trial 1–52), only head rotation data during the main experiment (trial 53–156) are analyzed in the present study.

For data analysis, it was necessary to match the timing information of the inertial sensor with events in the experiment. To this end, a time stamp was saved at the onset of the spatial cue in each trial and matched with the closest time stamp of the inertial sensor. Motion data were then epoched in the time interval –1 to +3s around spatial cue onset. For technical reasons, motion data were lost in some trials (up to 10 trials in 4/21 adults; up to 22 trials in 13 children).

To quantify horizontal head rotation to the left vs. right side, single-trial rotation around the yaw axis was extracted and averaged across trials, separately for experimental conditions. Other rotational axes (pitch & roll) were not of main interest in the present study. To calculate head orientation, angular velocity was integrated over time, using the function *cumtrapz* in Matlab.

### Task-dependent and -independent head rotation

We adapted an approach used in previous animal studies (Musall et al., 2019; Yin et al., 2025) to separate task-dependent from task-independent movement. In essence, task-dependent head rotation in the present study refers to head rotation explained by the task, i.e., by the spatial cue to the left or right in trials with lateral targets. For each individual participant, a series of linear models was used to regress head rotation individually at each time point (sampled from –1 to +2.95 s in steps of .05s) on the cue direction (left vs. right). Task-dependent movement (TDM) was quantified as the model prediction for select-left and select-right trials. To quantify TDM in the direction of the target (irrespective of cue direction), TDM for select-left trials was multiplied by –1, followed by averaging across cue directions. Task-independent head rotation (TIM) was quantified by calculating single-trial absolute deviation from the model prediction, followed by averaging across trials, which results in mean absolute error.

### Statistical analyses

Statistical analyses were performed in Matlab (R2023b) and Jamovi (Version 2.6.26.0). For behavioral data analyses, we used a one-way ANOVA on SNR with the factor age group (adults, younger children, older children), as well as mixed repeated-measures ANOVAs on the proportion of correct responses and response times with the additional factors: attention (selection, suppression) and side (left, right). Note that proportion data were transformed to rationalized arcsine units (Studebaker, 1985) to better meet the assumption of normality. For follow-up tests between age groups, we used independent-samples *t*-tests. For the analysis of the proportion of stream confusions (i.e., trials wherein a participant reported the distractor instead of the target), we used Wilcoxon rank sum tests since the data deviated substantially from the normal distribution.

To test for significant head rotation across time, we used within-subject cluster-based permutation tests to contrast head rotation within groups between conditions (e.g., select-left versus select-right; Gerber, 2025, two-sided, 10,000 permutations). To test for age-effects on task-dependent and task-independent movement (TDM & TIM), we used between-subject cluster-based permutation tests to contrast adults with children (younger and older), followed by independent-samples *t*-tests to contrast mean cluster effects between age groups. To relate movement to single-trial accuracy, we used generalized linear mixed-effects models (GLME) with a binomial distribution and logit link function (using the *fitglme* function in Matlab). Model parameters were estimated using the Laplace approximation. Significance of fixed effects was assessed using analysis of variance on the fitted model (using the *anova* function in Matlab). To test for interaction effects, we included the interaction term in an additional model and compared it to the reduced model (using the *compare* function in Matlab).

As effect sizes, we report eta-squared (*η^2^*) for one-way ANOVAs and partial eta-squared (*η^2^_p_*) for ANOVAs including more than one factor. For pairwise comparisons using post-hoc tests, we report Cohen’s *d*. Effect sizes (Cohen’s *d*) for cluster tests were derived from the mean and standard deviation of the data in the time intervals of significant clusters. Given the absence of an agreed-upon effect size measure for individual regressors in generalized linear mixed models, partial eta-squared (*η^2^_p_*) was approximated from the *F*-statistic and associated degrees of freedom (Cohen, 1973; *η^2^_p_* = (*F* * df1) / (*F* * df1 + df2)).

## Results

To investigate how head movements accompany attention deployment during development, this study used an auditory spatial attention task (Fig. 1) to compare behavioral performance and patterns of head rotation between three age groups of younger children (6–8 years; *n* = 25), older children (9–10 years; *n* = 20), and adults (18–30 years; *n* = 21).

### Behavioral task performance

To equalize overall task performance across participants to ∼71% accuracy, the signal-to-noise ratio (SNR) of target and distractor numbers was titrated per individual, using an adaptive staircase procedure (Fig. 1D; Levitt, 1971). An ANOVA revealed a significant effect of age group on SNR (Fig. 2A; *F_2,c3_*= 13.124; *p* < .001; *η^2^* = 0.294). The SNR was significantly lower for adults (*M_SNR_* = –28.6, *SE_SNR_* = 0.926) compared with older children (*M_SNR_* = – 24.95, *SE_SNR_* = 1.225; *t_39_* = –2.393, *p* = .0216; *d* = 0.748) and younger children (*M_SNR_* = –21.08, *SE_SNR_* = 1.021; *t_44_* = –5.367, *p* < .001; *d* = 1.589). Furthermore, older children reached lower SNR values than younger children (*t_43_* = –2.446, *p* = .019; *d* = 0.734). Thus, to ensure comparable performance levels, the acoustic scenario became on average less favorable (lower SNR) with increasing age.

**Figure 2.**
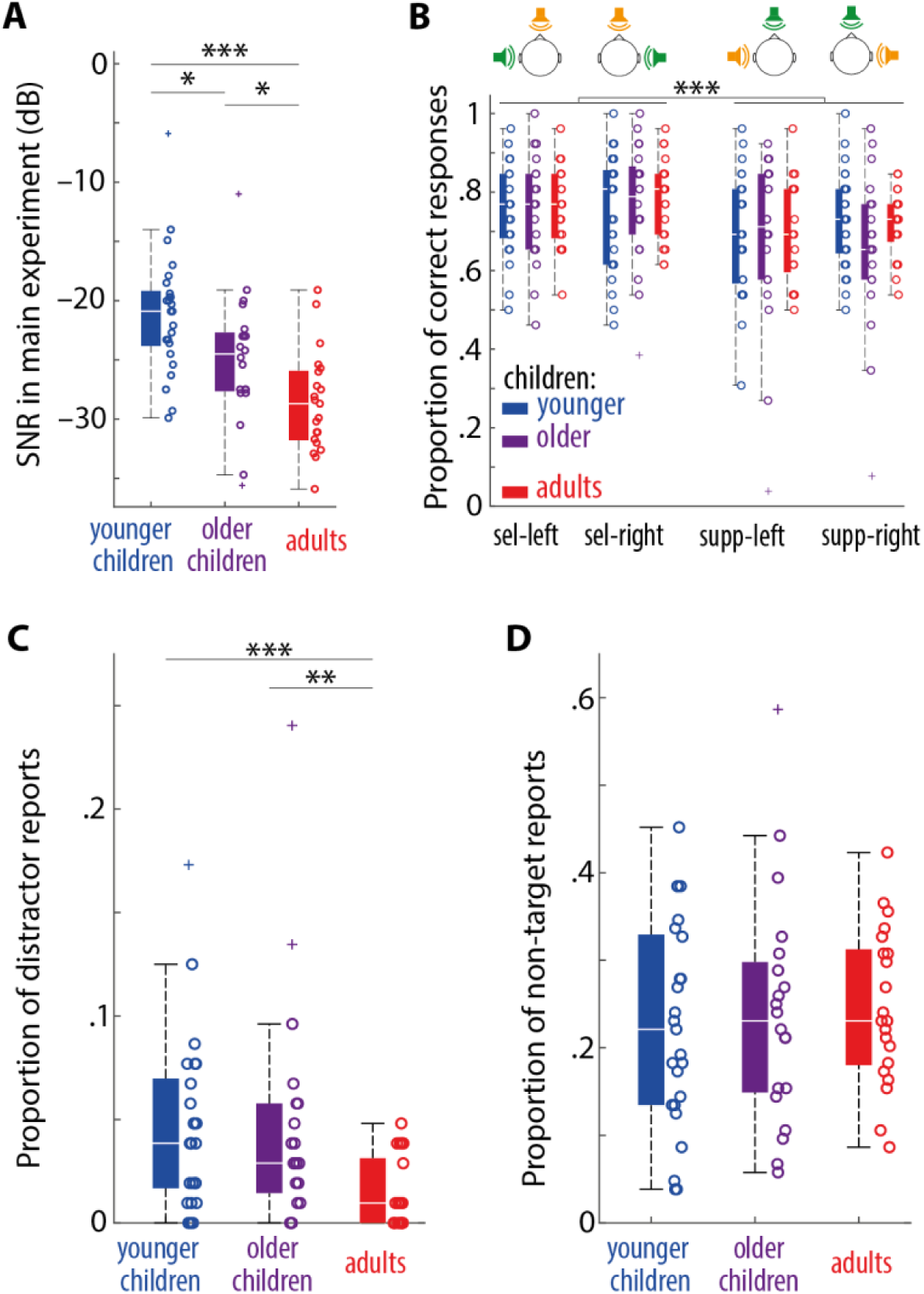
(**A**) Boxplots show the titrated SNR (used during the main experiment) as a function of age group. Circles show individual participants, crosses indicate outliers. (**B**) Boxplots show proportion of correct responses for different experimental conditions as a function of age group. Error trials were divided in “distractor report” (**C**) and “non-target report” (**D**; i.e., report of a number that was neither the target nor the distractor). Boxplots show respective proportions split by age group. * *p* < .05; ** *p* < .01; *** *p* < .001.

To analyze differences in task performance across experimental conditions, rau-transformed proportion correct scores were submitted to a mixed ANOVA with the within-subject factors attention (select, suppress) and side (left, right) as well as the between-subject factor age group (adults, young children, older children). The main effect age group and all interactions including age group were not significant (all *F* < 1.1; all *p* > .35), indicating that the individual titration of the SNR successfully resulted in comparable task accuracy across age groups. A significant main of effect of attention (Fig. 2B; *F_1, c3_* = 13.152; *p* < .001; *η^2^_p_ =* 0.173) revealed that accuracy was higher in trials with lateral versus frontal targets, which agrees with previous results in adult participants (Wöstmann et al., 2019). The main effect of side (*F_1, c3_* = 1.042; *p* = .311; *η^2^_p_ =* 0.016) and the attention x side interaction were not significant (*F_1, c3_* = .045; *p* = .832; *η^2^_p_ =* .001), indicating that task accuracy did not differ for left-versus right-sided presentation of the target number.

Although not of primary interest for this study, the same mixed ANOVA was used to analyze median response times (RT; Fig. S1). While none of the within-subject factors (attention, side) modulated RT (all *F* < 3.2, all *p* > .08), the effect of the between-subject factor age group was statistically significant (*F_2, 63_* = 30.766; *p* < .001; *η^2^_p_ =* 0.494), revealing shorter RTs in adults compared with younger children (*t_44_* = –7.112; *p* < .001; *d* = 2.105) and older children (*t_39_* = –7.828; *p* < .001; *d* = 2.446). RTs did not differ significantly in older versus younger children (*t_39_* = –1.647; *p* = .107; *d* = 0.494).

Importantly, to compare attentional control between age groups, we calculated the proportion of distractor reports, that is, trials wherein participants erroneously reported the distractor instead of the target (Fig. 2C). An ANOVA revealed a significant main effect of age group on rau-transformed proportions of distractor reports (*F_2, 63_* = 6.772; *p* = .002; *η^2^* = .177). As the proportion of distractor reports deviated from the normal distribution (especially so in the adult group), we also ran a non-parametric Kruskal-Wallis test, which confirmed the significant effect of age group (*χ^2^*(2) = 12.31; *p* = .002). Follow-up Wilcoxon rank sum tests revealed that adults reported fewer distractors than younger children (*Z* = –3.317; *p* < .001; *d* = .985) and older children (*Z* = –2.698; *p* = .007; *d* = .822). The number of distractor reports did not differ significantly between older and younger children (*Z* = –.344; *p* = .73; *d* = .038). Contrary to distractor reports, the proportion of non-target reports (i.e., reporting a number that was neither the target nor the distractor) did not differ between age groups (Fig. 2D, right; *F*_2,63_ = 0.38; *p* = .687; *η^2^* = .012). This demonstrates that despite overall comparable performance, target-distractor-confusions were more frequent in children.

### Stable patterns of average head rotation during development

We asked whether attention development would be accompanied by a systematic change of average head rotation patterns during auditory spatial attention. To our surprise, cluster-permutation tests revealed that spatial cues indicating upcoming target sounds on the left or right side induced remarkably similar average patterns of head rotation in the direction of the target in younger children (Fig. 3A; cluster *p*-value < .001, 0.65–2.2 s, *d* = 0.741), in older children (Fig. 3B; cluster *p*-value = .003, 0.55–2 s, *d* = 0.658), and in adults (Fig. 3C; cluster *p*-value < .001, 0.5–2 s, *d* = 0.882). Cue-induced head rotation in the time interval of the overlap of the three significant clusters shown in Figure 3A–C (0.65–2s) did not differ significantly between age groups (Fig 3D): average head rotation was submitted to a mixed ANOVA with the factors target side and age group, which did not reveal a significant main effect of age group (*F_2, 63_* = 0.682; *p* = .51; *η^2^* = 0.021) nor a significant age group x side interaction (*F_2, 63_* = 0.167; *p* = .846; *η^2^* = 0.005).

**Figure 3.**
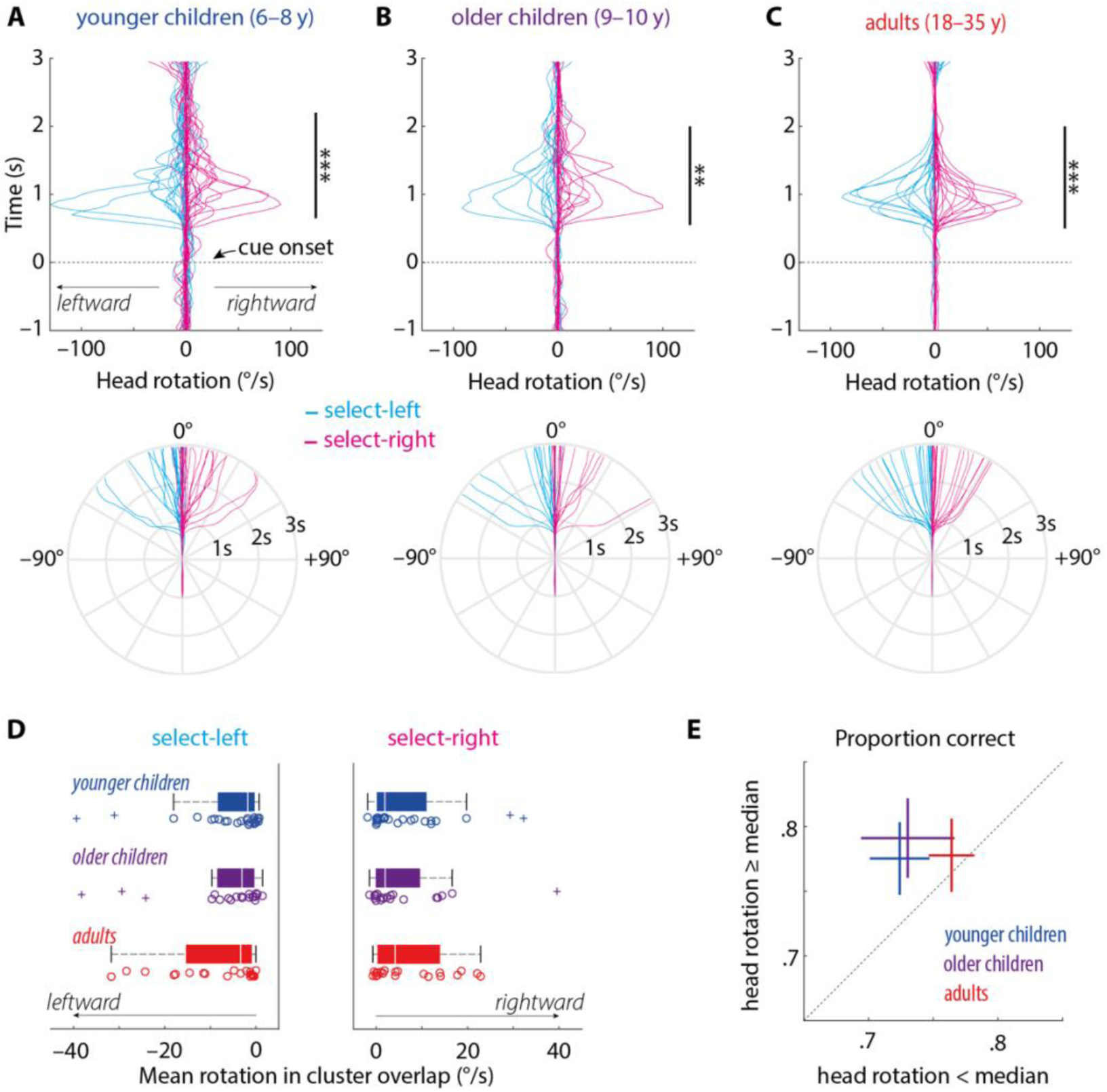
Line graphs (top) and polar plots (bottom) show respective angular velocity (in °/s) and head orientation (in °) as a function of time for younger children (**A**), older children (**B**), and adults (**C**). Each line represents the average of a single subject across trials in the select-left (blue) and select-right condition (pink). Vertical black bars indicate significant clusters; ** *p* < .01; *** *p* < .001. (**D**) Boxplots show mean head rotation in the time interval of cluster overlap (0.65–2 s) as a function of age group. Circles show individual participants, crosses indicate outliers. (**E**) The 45-degree plot contrasts proportion correct task performance for trials with head rotation below (x-axis) versus above the single-subject median (y-axis). Crosses show the mean ±1 SEM. Crosses above the diagonal display that higher proportion of correct responses was associated with stronger target-directed head rotation.

Spatial cueing of frontal targets (and thus implicit cueing of to-be-suppressed distraction on the left or right side) in suppress-left and suppress-right trials did not induce significant head rotation in children or adults (Fig. S2; all cluster *p-values* > .35), which agrees with a previous investigation in adults (Wöstmann & Obleser, 2026). For all further analyses, we thus focus on trials with lateral targets.

To test the relation of head rotation to lateral targets and accuracy, we used a GLME to regress single-trial accuracy on target-directed head rotation in the time interval of cluster overlap, target side, age group and a random subject intercept. A significant effect of head rotation amplitude on accuracy was found (*F_1,_*_3320_ = 4.55; *p* = .033; *η^2^_p_ =* .001). Including the head rotation x age group interaction term in the model did not improve model fit (*χ^2^*(2) = 2.197; *p* = .333; *ΔAIC* = +1.8, *ΔBIC* = +14.0). For visualization, we performed a median split on cue-induced head rotation, dividing trials into high- and low-rotation in the time interval of cluster overlap (Fig. 3E). The median split analysis shows that higher accuracy was associated with stronger target-directed head rotation.

This speaks to the functional relevance of target-directed head rotation for the success of auditory spatial attention. None of the other main effects were significant (all *p* > .39).

### Task-independent movement relates to attention development

Next, we tested whether variability of head rotation that could not be explained by the attention task would signify attention development. Contrary to average head rotation, trial-by-trial variability of head rotation decreased considerably from younger to older children and adults (Fig. 4A). To precisely quantify the variability in head rotation unrelated to the task instruction, we isolated single-trial task-independent from trial-average task-dependent movement. Task-dependent movement (TDM) was quantified as the model prediction resulting from a linear regression of head rotation on cue direction (left vs. right). Task-independent movement (TIM) was quantified as the mean absolute deviation of single-trial movement from the model prediction (Fig. 4B).

**Figure 4.**
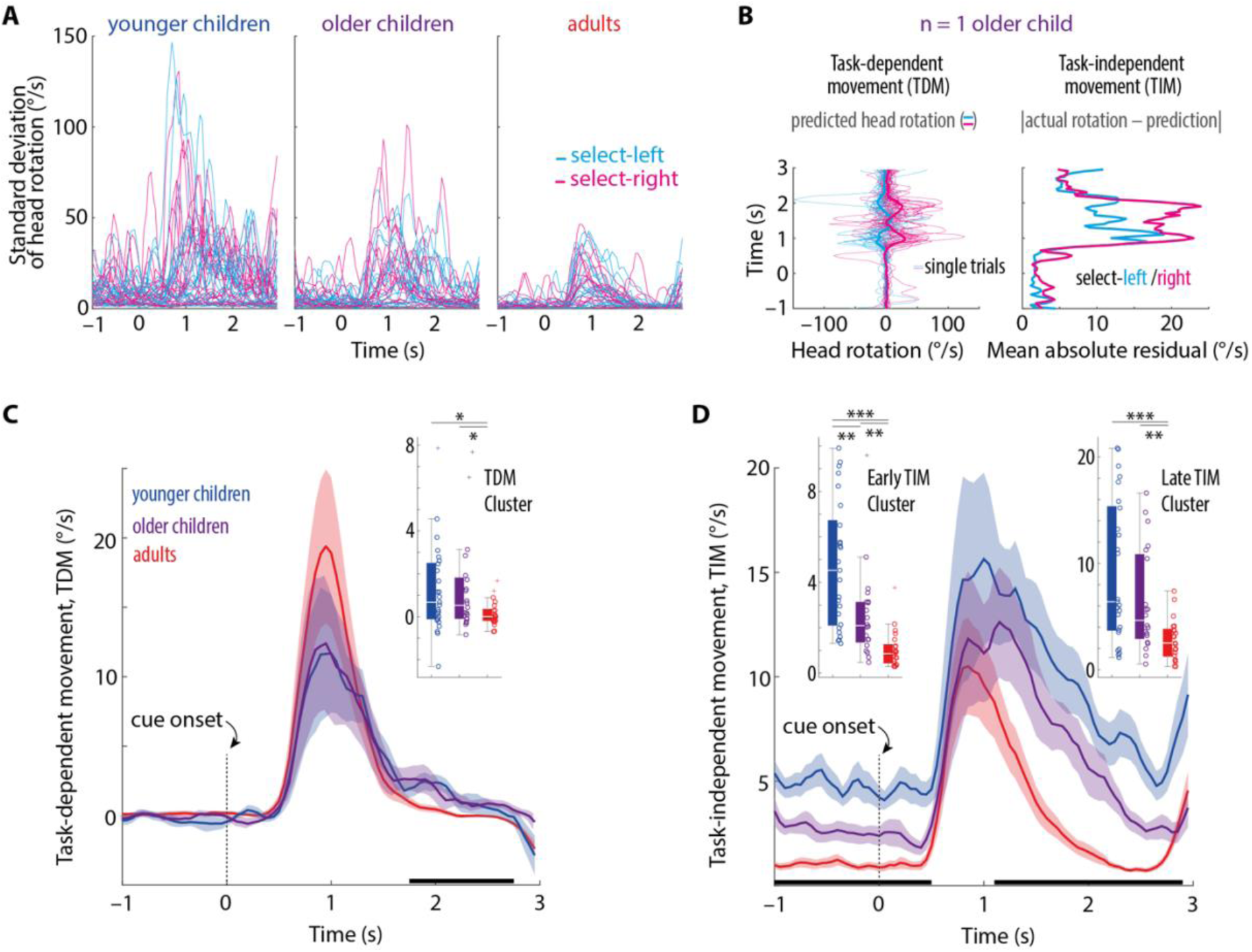
(**A**) Lines show the standard deviation of head rotation across trials for each individual participant in the three age groups, separately for select-left (blue) and select-right trials (pink). (**B**) Illustration of the approach to distinguish task-dependent movement (TDM) from task-independent movement (TIM). The figure shows head rotation data for one exemplary participant (n = 1 older child). TDM was estimated as the predicted head rotation resulting from a linear model to regress head rotation on cue direction (left vs. right). TIM was quantified as the absolute residual, that is, the unsigned deviation of single-trial rotation from the model prediction. Note that TDM is calculated for groups of trials per individual (select-left, select-right), whereas TIM is defined on the single-trial level (see single-trial GLME analysis reported in main text) but can be average across trials per condition (as depicted here). (**C**) Lines and error bands show mean ±1 SEM TDM (marginalized across select-left and select-right conditions), separated by age group. Horizontal black bars indicate significant clusters for the contrast of adults versus children (younger and older). Boxplots in the inset show mean TDM in the time interval of the respective cluster as a function of age group. Circles show individual participants, crosses indicate outliers. (**D**) Same as C but for TIM. * p < .05; ** *p* < .01; *** *p* < .001.

In line with our previous analysis (Fig. 3), task-dependent movement differed only slightly between age groups: In a significant cluster from 1.75 to 2.75s (found by contrasting TDM between children and adults), TDM was significantly lower in adults compared with older (Fig. 4C; *t*_39_ = – 2.354; *p* = .024; *d* = .735) and younger children (*t*_44_ = –2.346; *p* = .024; *d* = .695) but did not differ between younger and older children (*t*_43_ = 0.124; *p* = .902; *d* = .037). This indicates that children rotated their heads slightly more during a relatively late time interval in the trial just before the onset of competing sounds, presumably when adults already had reached their intended lateral orientation of the head.

Contrary, mean task-independent movement across trials differed markedly between age groups in two time intervals: In an early cluster mainly spanning the time interval before trial onset (Fig. 4D; –1 to +0.5s), TIM in adults was lower than in older (*t*_39_ = –3.23; *p* = .003; *d* = 1.008) and younger children (*t*_44_ = –5.891; *p* < .001; *d* = 1.744). Also, TIM in this early cluster was lower in older versus younger children (*t*_43_ = –2.931; *p* = .005; *d* = .879). Thus, differences in task-independent movement in the early cluster reflected a continuous developmental trajectory across childhood and into adulthood. Furthermore, TIM in a later cluster during the trial (1.1 to 2.9s) was significantly lower in adults versus older (*t*_39_ = –3.145; *p* = .003; *d* = .983) and younger children (*t*_44_ = –4.218; *p* < .001; *d* = 1.249), but did not differ between younger and older children (*t*_43_ = 1.56, *p* = .126; *d* = .468).

Finally, to test the relation of age-varying TIM and task performance, we used a GLME to regress single-trial accuracy on single-trial task-independent head rotation in the early and late TIM clusters and age group. We found a significant negative relation of task-independent head rotation with accuracy in the early cluster (–1 to +0.5s; *F_1,3320_* = 5.032; *p* = .025; *η^2^_p_ =* .002) but not in the late cluster (+1.1 to +2.9s; *F_1,3320_* < .01; *p* = .948; *η^2^_p_ <* .001). The main effect age group was not significant (*F_2,_*_3320_ = .02; *p* = .976; *η^2^_p_ <* .001). Including the TIM (early cluster) x age group interaction term in the model did not improve model fit (*χ^2^*(2) = .946; *p* = .623; *ΔAIC* = +3.1, *ΔBIC* = +15.2). For visualization, we performed a median split of TIM per individual participant, dividing trials into high- and low-TIM in the early cluster and assessed differences in task accuracy (Fig. 5). The median split analysis shows that higher accuracy was associated with less task-independent movement in the early TIM cluster. Together, these results show that task-independent movement before and early during a trial signifies both, between-subject differences in development and within-subject attentional mechanisms relevant for successful task performance.

**Figure 5.**
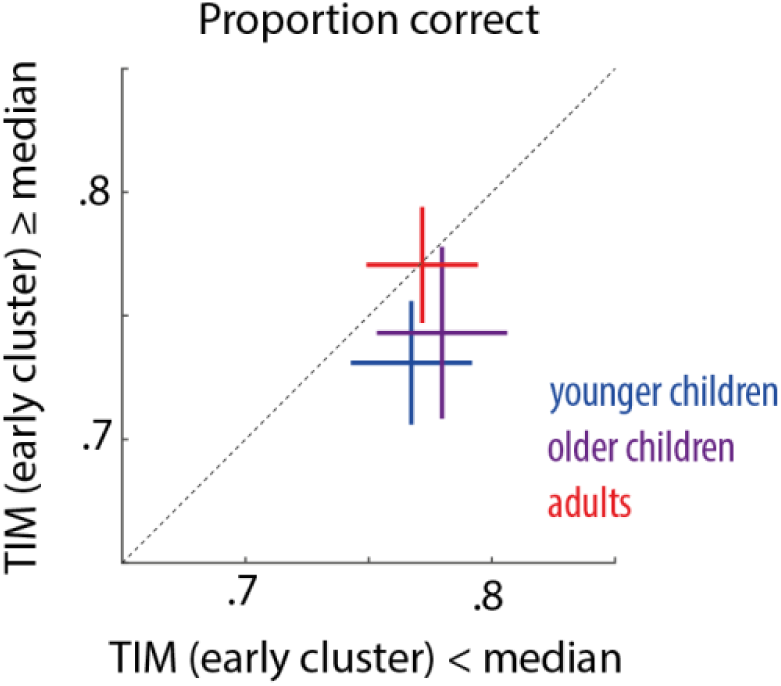
The 45-degree plot contrasts proportion correct task performance for trials with TIM in the early cluster below (x-axis) versus above the single-subject median (y-axis). Crosses show the mean ±1 SEM. Crosses below the diagonal indicate higher task accuracy for trials with less task-independent head rotation before and early during a trial (early TIM cluster; –1 to +0.5s).

## Discussion

The goal of the present study was to test whether the development of selective attention reflects in patterns of head movement. Surprisingly, we found that despite reduced attentional control in children (6–10 y) compared with adults, average patterns of head rotation in preparation for a lateral target sound were remarkably stable from childhood to adulthood. To the contrary, trial-by-trial deviation of head rotation from model-predicted movement (i.e., task-independent movement), decreased during development and explained task accuracy. Our results critically extend frameworks of embodied cognition (e.g., Clark, 1999; Wilson, 2002) and demonstrate that task-independent movement is not noise but an invaluable source of information to track selective attention and its development.

### Task-dependent head rotation is remarkably invariant to attention development

In a previous study using a similar paradigm, we found that young adult listeners performed prominent head rotations in the direction of lateral targets (Wöstmann & Obleser, 2026). Here, we show that 6–10-year-old children exhibit a strikingly similar pattern of results. Target-directed head rotations can support *binaural unmasking*, that is, enhanced sound detection and identification when competing sounds that arrive at the two ears differ acoustically (Culling & Lavandier, 2021; Kock, 1950). Head rotations of ∼30° are acoustically optimal when listening to a lateral target (90°) under distraction from the front (Jelfs et al., 2011). In the present study, the functional relevance of cue-induced head rotations was evidenced by higher task accuracy in trials with larger target-directed head rotation. But do acoustically beneficial head rotations induced by a spatial attention cue reflect development of selective attention?

Auditory developmental research has shown that attentional control is reduced in younger children compared with adults and increases during middle childhood (∼6–10 years), reflecting in decreasing distractibility (Calcus, 2024; Hoyer et al., 2021; Wetzel et al., 2019). In agreement, we found a higher number of distractor reports in children compared with adults. If cue-induced head rotation to a lateral target constitutes an embodied mechanism of attention development, task-dependent head rotation should increase during childhood development as well. However, to our surprise, we found that patterns of task-dependent head rotation were largely stable across development. Only relatively late during a trial after the peak of average head rotation, we found stronger (instead of expected weaker) task-dependent head rotation in children versus adults, but no difference between younger and older children. This could reflect the still-maturing capacity to terminate motor engagement once its functional contribution has diminished, consistent with evidence for a protracted improvement in inhibitory control from childhood through adulthood (Williams et al., 1999).

Together, these results suggest that cue-induced head rotation does not reflect the development of attentional control after the age of 6 years. This developmental stability of a functionally beneficial motor strategy indicates that, rather than emerging alongside higher-order attentional control, embodied contributions to spatial hearing may be established early and recruited in a largely mature form throughout childhood, consistent with evidence that basic spatial listening skills are substantially developed before school age (Litovsky, 2005; Van Deun et al., 2009).

### Task-independent head rotation reflects development of spatial attention

Research in mice has shown that movements not explained by task variables (i.e., task-independent movement) strongly shape neural activity (Musall et al., 2019) and increase during disengagement (Yin et al., 2025). Here, we found that variability of head rotations in humans prominently decreased during childhood development, speaking to a developmental increase in target-directed movement stereotypy (see also Golenia et al., 2018).

The major research question of the present study was whether isolating task-independent from task-dependent movement would help to predict the development and success of spatial selective attention. Our results demonstrate temporal specificity of task-independent head rotation. Overall movement was strongest at ∼1s following spatial cue presentation, when both task-dependent and -independent movement peaked. Interestingly, age-effects on task-independent movement surrounded this overall peak in movement and were found in an earlier time window (covering the pre-trial baseline and cue presentation) and in a later time window (covering the delay period and parts of sound presentation). Arguably, task-independent movement in these time windows reflects distinct underlying processes.

Task-independent movement following cue presentation quantifies the deviation from average head rotation in the respective cue-condition and thus reflects inconsistency of goal-directed motor activity in service of spatial attention. We found recently that cue-induced head rotation in adults relates positively to a temporally preceding neural signature of spatial attention (i.e., lateralized alpha oscillations; Wöstmann & Obleser, 2026). This might suggest that head rotation variability after cue presentation reflects variability in the acting-out of attention. However, task-independent movement following cue presentation did not differ significantly between younger and older children did not relate to task accuracy, meaning that it does not reflect the development and success of selective attention.

In theory, optimal preparation for an upcoming spatial attention task with unknown assignment of target and distractor sounds to two locations is to remain still and to focus on the upcoming attention cue. Thus, any movement before and early during a trial likely reflects task disengagement (see also Seli et al., 2014). Our results suggest that disengagement-reflecting head rotation decreases continuously during development (from younger to older childhood and to adulthood) and that less disengagement-related head rotation is beneficial for attention success. This might suggest that learning to withhold task-irrelevant movement is itself part of the developmental trajectory of selective attention (Casey et al., 2005).

Beyond the theoretical value of task-independent movement for understanding the development of spatial attention, our findings suggest that it might serve as a real-time behavioral proxy for attentional engagement (e.g., in classroom or clinical settings), flagging moments or individuals at risk of reduced performance without interrupting the ongoing task. Monitoring the temporal structure and specifically the task-independent component of movement may offer a scalable, unobtrusive route toward identification of attentional difficulties and toward evaluating whether interventions and technologies successfully reduce disengagement-related movement to improve attention performance.

### Limitations

Individual adjustment of the acoustic SNR could contribute to observed differences in performance and movement between age groups. However, the rationale was to control for audibility and overall task performance to contrast listeners from different age groups in a task wherein they all reached comparable levels of accuracy (McCreery & Stelmachowicz, 2011). Gyroscopic tracking with a sensor placed on the head does not allow to differentiate between some movement types, such as rotation of the head alone versus rotation of the upper body (including the head). However, our study was designed to primarily elicit horizontal rotations of the head. Nevertheless, future studies should use more comprehensive approaches to motion tracking to improve our understanding of the specificity of task-dependent and -independent movement for spatial attention and its development.

## Conclusion

This study shows that embodied signatures of cognitive development are not uniform. Development selectively refines the consistency of motor behavior, while the underlying orienting strategy is already adult-like in middle-childhood. Our results position movement as a measurable, developmentally meaningful component of attention. These findings extend embodied cognition frameworks to a developmental context and point to task-independent movement as an invaluable non-invasive marker of attentional maturation.

## Supplementary Information

**Figure S1.**
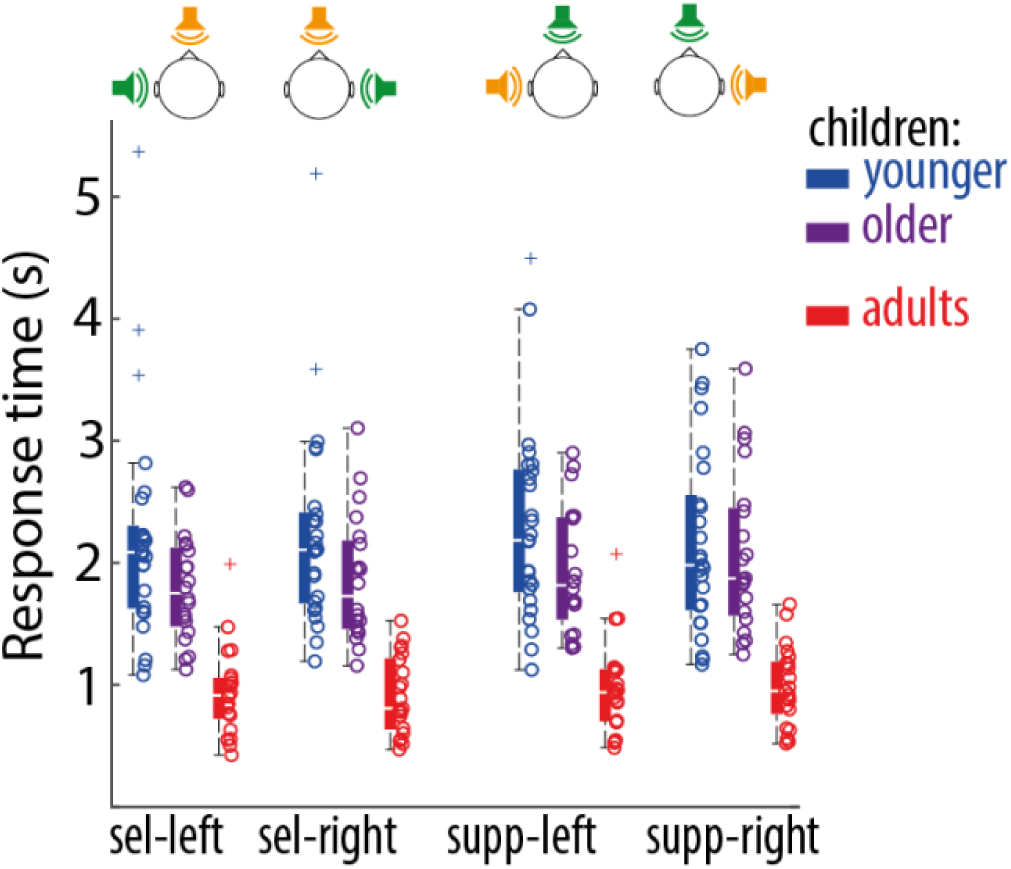
Boxplots show median response time for different experimental conditions as a function of age group. Circles show individual participants, crosses indicate outliers. Response times correspond to the time interval between presentation of the response screen (‘?’) and the button press.

**Figure S2.**
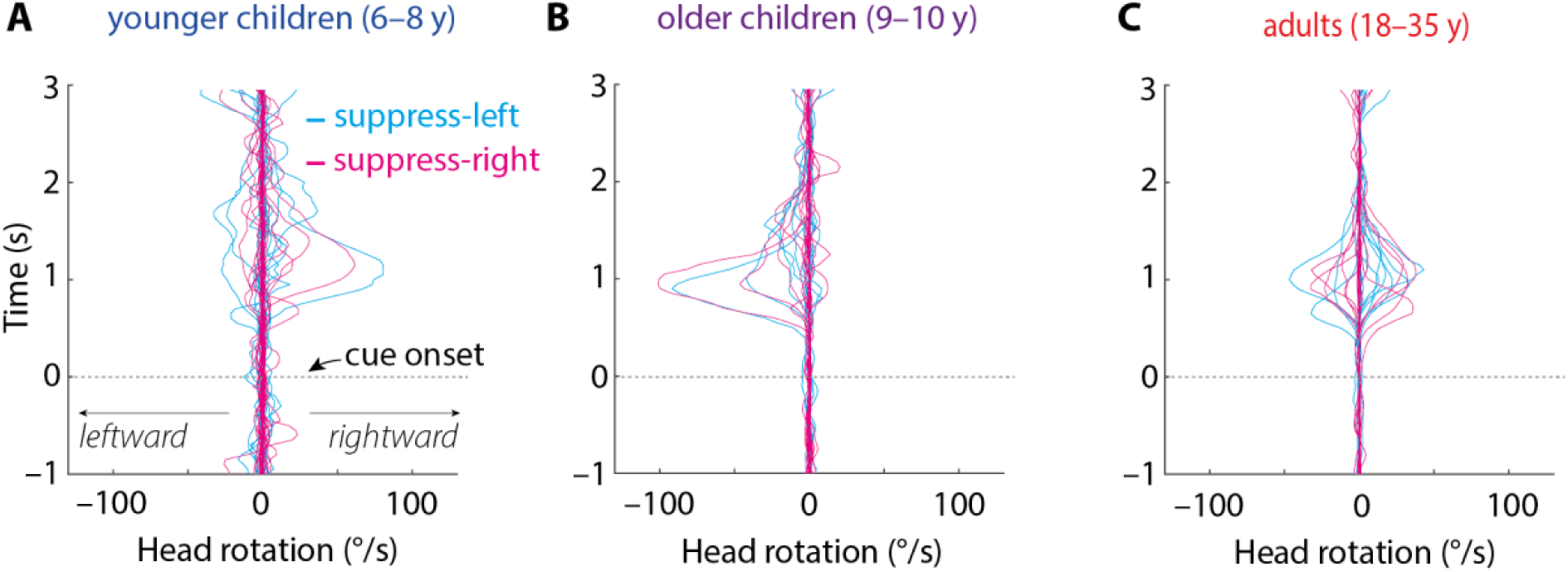
Line graphs show respective angular velocity (in °/s) as a function of time for younger children (**A**), older children (**B**), and adults (**C**). Each line represents the average of a single subject across trials in the suppress-left (cyan) and suppress-right condition (magenta).

## References

Agostinho, S., Tindall-Ford, S., Ginns, P., Howard, S. J., Leahy, W., & Paas, F. (2015). Giving Learning a Helping Hand: Finger Tracing of Temperature Graphs on an iPad. Educational Psychology Review, 27(3), 427–443. 10.1007/s10648-015-9315-5

Brainard, D. H. (1997). The Psychophysics Toolbox. Spatial Vision, 10(4), 433–436. 10.1163/156856897X00357

Calcus, A. (2024). Development of auditory scene analysis: A mini-review. Frontiers in Human Neuroscience, 18. 10.3389/fnhum.2024.1352247

Casey, B. J., Tottenham, N., Liston, C., & Durston, S. (2005). Imaging the developing brain: What have we learned about cognitive development? Trends in Cognitive Sciences, 9(3), 104–110. 10.1016/j.tics.2005.01.011

Caviola, S., Visentin, C., Borella, E., Mammarella, I., & Prodi, N. (2021). Out of the noise: Effects of sound environment on maths performance in middle-school students. Journal of Environmental Psychology, 73, 101552. 10.1016/j.jenvp.2021.101552

Cisek, P. (2007). Cortical mechanisms of action selection: The affordance competition hypothesis. Philosophical Transactions of the Royal Society B: Biological Sciences, 362(1485), 1585–1599. 10.1098/rstb.2007.2054

Clark, A. (1999). An embodied cognitive science? Trends in Cognitive Sciences, 3(9), 345–351. 10.1016/S1364-6613(99)01361-3

Cohen, J. (1973). Eta-Squared and Partial Eta-Squared in Fixed Factor Anova Designs. Educational and Psychological Measurement, 33(1), 107–112. 10.1177/001316447303300111

Criscuolo, A., Schwartze, M., & Kotz, S. A. (2022). Cognition through the lens of a body–brain dynamic system. Trends in Neurosciences, 45(9), 667–677. 10.1016/j.tins.2022.06.004

Culling, J. F., & Lavandier, M. (2021). Binaural Unmasking and Spatial Release from Masking. In R. Y. Litovsky, M. J. Goupell, R. R. Fay, & A. N. Popper (Eds.), Binaural Hearing: With 93 Illustrations (pp. 209–241). Springer International Publishing. 10.1007/978-3-030-57100-9_8

Desimone, R., & Duncan, J. (1995). Neural mechanisms of selective visual attention. Annual Review of Neuroscience, 18, 193–222. 10.1146/annurev.ne.18.030195.001205

Dockrell, J. E., & Shield, B. M. (2006). Acoustical barriers in classrooms: The impact of noise on performance in the classroom. British Educational Research Journal, 32(3), 509–525. 10.1080/01411920600635494

Eaton, W. O., McKeen, N. A., & Campbell, D. W. (2001). The Waxing and Waning of Movement: Implications for Psychological Development. Developmental Review, 21(2), 205–223. 10.1006/drev.2000.0519

Gerber, E. M. (2025). Permutest (Version https://www.mathworks.com/matlabcentral/fileexchange/71737-permutest) [Computer software]. MATLAB Central File Exchange. https://www.mathworks.com/matlabcentral/fileexchange/71737-permutest

Golenia, L., Schoemaker, M. M., Otten, E., Mouton, L. J., & Bongers, R. M. (2018). Development of reaching during mid-childhood from a Developmental Systems perspective. PLOS ONE, 13(2), e0193463. 10.1371/journal.pone.0193463

Grange, J. A., & Culling, J. F. (2016). The benefit of head orientation to speech intelligibility in noise. The Journal of the Acoustical Society of America, 139(2), Article 2. 10.1121/1.4941655

Hoyer, R. S., Elshafei, H., Hemmerlin, J., Bouet, R., & Bidet-Caulet, A. (2021). Why Are Children So Distractible? Development of Attention and Motor Control From Childhood to Adulthood. Child Development, 92(4), e716–e737. 10.1111/cdev.13561

Janssen, M., Chinapaw, M. J. M., Rauh, S. P., Toussaint, H. M., van Mechelen, W., & Verhagen, E. A. L. M. (2014). A short physical activity break from cognitive tasks increases selective attention in primary school children aged 10–11. Mental Health and Physical Activity, 7(3), 129–134. 10.1016/j.mhpa.2014.07.001

Jelfs, S., Culling, J. F., & Lavandier, M. (2011). Revision and validation of a binaural model for speech intelligibility in noise. Hearing Research, 275(1), Article 1. 10.1016/j.heares.2010.12.005

Kock, W. E. (1950). Binaural Localization and Masking. The Journal of the Acoustical Society of America, 22(6), Article 6. (world). 10.1121/1.1906692

Kondo, H. M., Pressnitzer, D., Toshima, I., & Kashino, M. (2012). Effects of self-motion on auditory scene analysis. Proceedings of the National Academy of Sciences, 109(17), Article 17. 10.1073/pnas.1112852109

Ladouce, S., Donaldson, D. I., Dudchenko, P. A., & Ietswaart, M. (2017). Understanding Minds in Real-World Environments: Toward a Mobile Cognition Approach. Frontiers in Human Neuroscience, 10. 10.3389/fnhum.2016.00694

Levitt, H. (1971). Transformed Up-Down Methods in Psychoacoustics. The Journal of the Acoustical Society of America, 49(2B), 467–477. 10.1121/1.1912375

Litovsky, R. Y. (2005). Speech intelligibility and spatial release from masking in young children. The Journal of the Acoustical Society of America, 117(5), 3091–3099. 10.1121/1.1873913

Loeffler, J., Raab, M., & Cañal-Bruland, R. (2016). A Lifespan Perspective on Embodied Cognition. Frontiers in Psychology, 7. 10.3389/fpsyg.2016.00845

Mathis, A., Mamidanna, P., Cury, K. M., Abe, T., Murthy, V. N., Mathis, M. W., & Bethge, M. (2018). DeepLabCut: Markerless pose estimation of user-defined body parts with deep learning. Nature Neuroscience, 21(9), 1281–1289. 10.1038/s41593-018-0209-y

McCreery, R. W., & Stelmachowicz, P. G. (2011). Audibility-based predictions of speech recognition for children and adults with normal hearing. The Journal of the Acoustical Society of America, 130(6), 4070–4081. 10.1121/1.3658476

Morillon, B., Hackett, T. A., Kajikawa, Y., & Schroeder, C. E. (2015). Predictive motor control of sensory dynamics in auditory active sensing. Current Opinion in Neurobiology, SI: Brain Rhythms and Dynamic Coordination, 31, 230–238. 10.1016/j.conb.2014.12.005

Musall, S., Kaufman, M. T., Juavinett, A. L., Gluf, S., & Churchland, A. K. (2019). Single-trial neural dynamics are dominated by richly varied movements. Nature Neuroscience, 22(10), 1677–1686. 10.1038/s41593-019-0502-4

Noonan, M. P., Crittenden, B. M., Jensen, O., & Stokes, M. G. (2018). Selective inhibition of distracting input. Behavioural Brain Research, 355, 36–47. 10.1016/j.bbr.2017.10.010

Oldfield, R. C. (1971). The assessment and analysis of handedness: The Edinburgh inventory. Neuropsychologia, 9(1), Article 1. 10.1016/0028-3932(71)90067-4

Pereira, T. D., Shaevitz, J. W., & Murthy, M. (2020). Quantifying behavior to understand the brain. Nature Neuroscience, 23(12), 1537–1549. 10.1038/s41593-020-00734-z

Ping, R., & Goldin-Meadow, S. (2010). Gesturing Saves Cognitive Resources When Talking About Nonpresent Objects. Cognitive Science, 34(4), 602–619. 10.1111/j.1551-6709.2010.01102.x

Prins, N., & Kingdom, F. A. A. (2018). Applying the Model-Comparison Approach to Test Specific Research Hypotheses in Psychophysical Research Using the Palamedes Toolbox. Frontiers in Psychology, S. 10.3389/fpsyg.2018.01250

Rapport, M. D., Bolden, J., Kofler, M. J., Sarver, D. E., Raiker, J. S., & Alderson, R. M. (2009). Hyperactivity in Boys with Attention-Deficit/Hyperactivity Disorder (ADHD): A Ubiquitous Core Symptom or Manifestation of Working Memory Deficits? Journal of Abnormal Child Psychology, 37(4), 521–534. 10.1007/s10802-008-9287-8

Reigal, R. E., Barrero, S., Martín, I., Morales-Sánchez, V., Juárez-Ruiz de Mier, R., C Hernández-Mendo, A. (2019). Relationships Between Reaction Time, Selective Attention, Physical Activity, and Physical Fitness in Children. Frontiers in Psychology, 10. 10.3389/fpsyg.2019.02278

Rizzolatti, G., Riggio, L., Dascola, I., C Umiltá, C. (1987). Reorienting attention across the horizontal and vertical meridians: Evidence in favor of a premotor theory of attention. Neuropsychologia, 25(1, Part 1), 31–40. 10.1016/0028-3932(87)90041-8

Schroeder, C. E., Wilson, D. A., Radman, T., Scharfman, H., & Lakatos, P. (2010). Dynamics of Active Sensing and perceptual selection. Current Opinion in Neurobiology, Cognitive Neuroscience, 20(2), Article 2. 10.1016/j.conb.2010.02.010

Seli, P., Carriere, J. S. A., Thomson, D. R., Cheyne, J. A., Martens, K. A. E., & Smilek, D. (2014). Restless mind, restless body. Journal of Experimental Psychology: Learning, Memory, and Cognition, 40(3), 660–668. 10.1037/a0035260

Stangl, M., Maoz, S. L., & Suthana, N. (2023). Mobile cognition: Imaging the human brain in the ‘real world’. Nature Reviews Neuroscience, 24(6), 347–362. 10.1038/s41583-023-00692-y

Stringer, C., Pachitariu, M., Steinmetz, N., Reddy, C. B., Carandini, M., & Harris, K. D. (2019). Spontaneous behaviors drive multidimensional, brainwide activity. Science, 364(6437), eaav7893. 10.1126/science.aav7893

Studebaker, G. A. (1985). A ‘Rationalized’ Arcsine Transform. Journal of Speech, Language, and Hearing Research, 28(3), 455–462. 10.1044/jshr.2803.455

Van Deun, L., van Wieringen, A., Van den Bogaert, T., Scherf, F., Offeciers, F. E., Van de Heyning, P. H., Desloovere, C., Dhooge, I. J., Deggouj, N., De Raeve, L., & Wouters, J. (2009). Sound localization, sound lateralization, and binaural masking level differences in young children with normal hearing. Ear and Hearing, 30(2), 178–190. 10.1097/AUD.0b013e318194256b

Wetzel, N., Scharf, F., & Widmann, A. (2019). Can’t Ignore—Distraction by Task-Irrelevant Sounds in Early and Middle Childhood. Child Development, 90(6), Article 6. 10.1111/cdev.13109

Williams, B. R., Ponesse, J. S., Schachar, R. J., Logan, G. D., & Tannock, R. (1999). Development of inhibitory control across the life span. Developmental Psychology, 35(1), 205–213. 10.1037/0012-1649.35.1.205

Wilson, M. (2002). Six views of embodied cognition. Psychonomic Bulletin & Review, 9(4), 625–636. 10.3758/BF03196322

Wöstmann, M., Alavash, M., & Obleser, J. (2019). Alpha oscillations in the human brain implement distractor suppression independent of target selection. The Journal of Neuroscience, 39(49), Article 49. 10.1523/JNEUROSCI.1954-19.2019

Wöstmann, M., Meineke, H. M., Schönweiler, R., Hollfelder, D., Bruchhage, K.-L., Leichtle, A., & Obleser, J. (2025). Neural alpha oscillations and auditory steady-state responses during adaptation to a cochlear implant. Cerebral Cortex, 35(8), bhaf244. 10.1093/cercor/bhaf244

Wöstmann, M., & Obleser, J. (2026). Spatial attention in the moving brain: Dissociable roles of neural alpha oscillations and head rotation. iScience, 29(9). 10.1016/j.isci.2026.117041

Wöstmann, M., Störmer, V. S., Obleser, J., Addleman, D. A., Andersen, Søren K., Gaspelin, N., Geng, J. J., Luck, S. J., Noonan, M. P., Slagter, H. A., & Theeuwes, J. (2022). Ten simple rules to study distractor suppression. Progress in Neurobiology, 213, 102269. 10.1016/j.pneurobio.2022.102269

Yin, C., Melin, M. D., Rojas-Bowe, G., Sun, X. R., Couto, J., Gluf, S., Kostiuk, A., Musall, S., & Churchland, A. K. (2025). Spontaneous movements and their relationship to neural activity fluctuate with latent engagement states. Neuron, 113(18), 3048–3063.e5. 10.1016/j.neuron.2025.06.001

